# GS-967 and Eleclazine Block Sodium Channels in Human Induced Pluripotent Stem Cell-derived Cardiomyocytes

**DOI:** 10.1101/2020.05.08.084350

**Authors:** Franck Potet, Defne E. Egecioglu, Paul W. Burridge, Alfred L. George

## Abstract

GS-967 and eleclazine (GS-6615) are novel sodium channel inhibitors exhibiting antiarrhythmic effects in various *in vitro* and *in vivo* models. The antiarrhythmic mechanism has been attributed to preferential suppression of late sodium current (*I*_NaL_). Here, we took advantage of a throughput automated electrophysiology platform (SyncroPatch 768PE) to investigate the molecular pharmacology of GS-967 and eleclazine on peak sodium current (*I*_NaP_) recorded from human induced pluripotent stem cell (hiPSC)-derived cardiomyocytes. We compared GS-967 and eleclazine to the antiarrhythmic drug lidocaine, the prototype *I*_NaL_ inhibitor ranolazine, and the slow inactivation enhancing drug lacosamide. In human induced pluripotent stem cell-derived cardiomyocytes, GS-967 and eleclazine caused a reduction of *I*_NaP_ in a frequency-dependent manner consistent with use-dependent block (UDB). GS-967 and eleclazine had similar efficacy but evoked more potent UDB of *I*_NaP_ (IC_50_=0.07 and 0.6 μM, respectively) than ranolazine (7.8 μM), lidocaine (133.5 μM) and lacosamide (158.5 μM). In addition, GS-967 and eleclazine exerted more potent effects on slow inactivation and recovery from inactivation compared to the other sodium channel blocking drugs we tested. The greater UDB potency of GS-967 and eleclazine was attributed to the significantly higher association rates (K_ON_) and moderate unbinding rate (K_OFF_) of these two compounds with sodium channels. We propose that substantial UDB contributes to the observed antiarrhythmic efficacy of GS-967 and eleclazine.

**SIGNIFICANCE STATEMENT:** We investigated the molecular pharmacology of GS-967 and eleclazine on sodium channels in human induced pluripotent stem cell derived cardiomyocytes using a high throughput automated electrophysiology platform. Sodium channel inhibition by GS-967 and eleclazine has unique features including accelerating the onset of slow inactivation and impairing recovery from inactivation. These effects combined with rapid binding and moderate unbinding kinetics explain potent use-dependent block, which we propose contributes to their observed antiarrhythmic efficacy.

## INTRODUCTION

Sodium current (*I*_Na_) in cardiac myocytes carried primarily by Na_V_1.5 channels is responsible for the rapid upstroke of atrial and ventricular action potentials as well as the rapid propagation of depolarization throughout the heart. Sodium channels can transition between at least three distinct states: resting (closed), open (active) and inactivated (non-conducting) (Hodgkin and Huxley, 1952). The transitions between these states are voltage and time dependent. Antiarrhythmic, anticonvulsant and local anesthetic agents have been shown to block the propagation of action potentials by interacting differently with each of the states of the channel (Strichartz et al., 1987). In cardiac tissue, the main effect of antiarrhythmic drugs is to prevent abnormal electrical impulse propagation and conduction, thus suppressing non-pacemaker generated electrical activity arising from damaged cardiac myocytes.

Effective class I antiarrhythmic drugs such as lidocaine, which block voltage-gated sodium channels, exhibit greater efficacy in situations associated with rapid repetitive firing of action potentials (use-dependent block) or prolonged tissue depolarization, such as in myocardial ischemia. These drugs should not affect normal cellular excitability. Thus, effective arrhythmia suppression depends on the properties of the drug molecule that convey high affinity binding to the channel pore when the channel is in the open or inactivated state. Such high affinity binding results in a slowed recovery of the drug-bound channel from inactivation as the cell membrane repolarizes (Ragsdale et al., 1996).

Disturbances in Na_V_1.5 function can promote life-threatening cardiac arrhythmia. When Na_V_1.5 fails to inactivate fully after opening, Na^+^ influx continues throughout the action potential plateau. The resulting current, referred as late *I*_Na_ (*I*_NaL_), can promote prolongation of the action potential duration. Late *I*_Na_ is normally small, but its amplitude is greater in certain acquired or heritable conditions, including failing and/or ischemic heart (Le Grand et al., 1995), oxidative stress (Song et al., 2006), or mutations in *SCN5A*, which encodes Na_V_1.5 (Bennett et al., 1995; Ruan et al., 2009). *SCN5A* mutations that cause enhanced *I*_NaL_ produce type 3 long QT syndrome (LQT3) (Antzelevitch et al., 2014). Drugs that selectively suppress *I*_NaL_ may offer a targeted antiarrhythmic strategy in these conditions.

Many local anesthetic and antiarrhythmic agents have greater potency to block *I*_NaL_ than peak *I*_Na_ (*I*_NaP_). Certain compounds such as ranolazine (Gupta et al., 2015) and F15845 (Pignier et al., 2010), are described as preferential *I*_NaL_ blockers. GS-967 (a triazolopyridine derivative, 6-(4-(trifluoromethoxy)phenyl)-3-(trifluoromethyl)-[1,2,4]triazolo[4,3-a]pyridine also referred as PRAX-330) (Koltun et al., 2016) and eleclazine (dihydrobenzoxazepinone, formerly known as GS-6615) are recently described sodium channel blockers that were originally demonstrated to exert potent antiarrhythmic effects in rabbit ventricular, canine and pig atrial myocytes by a proposed mechanism of action involving preferential *I*_NaL_ block (Belardinelli et al., 2013; Fuller et al., 2016; Sicouri et al., 2013). We previously demonstrated that GS-967, in addition to blocking *I*_*NaL*_, also exerts a strong use-dependent block (UDB) of *I*_NaP_ conducted by heterologously expressed recombinant human Na_V_1.5 and proposed that this phenomenon might contribute to its antiarrhythmic effect (Potet et al., 2016). Further studies examining the molecular pharmacology of GS-967 and eleclazine on sodium channels in cardiomyocytes should provide valuable insight into the drugs’ mechanism of action in a therapeutically relevant environment.

In this study, we investigated the molecular pharmacology of GS-967 and eleclazine in hiPSC-derived cardiomyocytes using a high throughput automated electrophysiology platform (SyncroPatch 768PE). We compared GS-967 and eleclazine to lidocaine (a class 1-b antiarrhythmic drug that promotes UDB) (Herzog et al., 2003), to ranolazine (a prototype *I*_*NaL*_ inhibitor with class1-b antiarrhythmic characteristics) (Szel et al., 2011), and lacosamide (an anticonvulsant drug that enhances slow inactivation) (Errington et al., 2008). Both, ranolazine and lacosamide can enhance slow inactivation of Na_V_ channels (Errington et al., 2008; Kahlig et al., 2014). We observed that eleclazine, like GS-967, exhibit moderate dissociation rates, comparable to the class 1-b antiarrhythmic lidocaine, along with uniquely rapid binding kinetics. These properties explain the potent use-dependent block of *I*_NaP_ by GS-967 and eleclazine observed in hiPSC-derived cardiomyocytes.

## MATERIALS AND METHODS

### Cell Culture

Human iPSC were derived from peripheral blood of a male donor as previously described (Burridge et al., 2016). The resulting line was designated 113c4. The hiPSC line was cultured on growth factor-reduced Matrigel (Corning, NY, USA) in E8 medium (Burridge et al., 2015). Cardiac differentiation was completed in CDM3 as previously described (Burridge et al., 2014). CDM3 consisted of RPMI 1640 (Corning), 500 μg/ml fatty acid-free albumin (GenDEPOT, Barker, TX, USA), and 200 μg/ml L-ascorbic acid 2-phosphate (Wako, Richmond, VA, USA). At day 20 (d20) of differentiation, cells were dissociated by incubation in DPBS (without Mg^2+^ or Ca^2+^) for 20 min at 37°C and then in 1:200 Liberase TH (Roche, Basel, Switzerland) in DPBS for 20 min at 37°C. Cell were collected by centrifuging at 200 ×*g* for 3 min, counted, and replated in Matrigel-coated 6-well plates at 2-4 million cells per well in CDM3 supplemented with 40% FBS (Opti-Gold, GenDEPOT). Cells were returned to CDM3 on d22. Human iPSC-derived cardiomyocytes were re-plated in 30 mm culture dishes 5 days prior to the experiment.

### Automated patch clamp recording

Automated patch clamp recording was performed using a Syncropatch 768 PE (Nanion Technologies, Munich, Germany). On the day of the experiment, cells were washed once with DPBS (Mg/Ca free) for 20 minutes. Cells were then detached with 5 min treatment with TrypLE followed by 20-30 minutes treatment with CDM3 media with 1:200 dilution of Liberase TH. Cells were then re-suspended in 15% CDM3 media and 85% external solution at 170,000 cells/ml. Cells were allowed to recover for at least 30 min at 15°C while shaking on a rotating platform. Following equilibration, 10 μl of cell suspension was added to each well of a 384-well, single-hole, low resistance (3 MΩ) NPC-384 ‘chip’ (Nanion Technologies).

Pulse generation and data collection were done with PatchController384 V.1.3.0 and DataController384 V1.2.1 (Nanion Technologies). Whole-cell currents were filtered at 3 kHz and acquired at 10 kHz. The access resistance and apparent membrane capacitance were estimated using protocols within the data acquisition software. Series resistance was compensated 95% and leak and capacitance artifacts were subtracted out using the P/4 method. Whole-cell currents were recorded at room temperature (∼ 24°C) in the whole-cell configuration. The external solution contained (in mM): NaCl 140, KCl 4, CaCl_2_ 2, MgCl_2_ 1, HEPES 10, glucose 5, pH = 7.4. The internal solution contained (in mM): CsF 110, CsCl 10, NaCl 10, HEPES 10, EGTA 20, pH = 7.2.

Using hiPSC-derived cardiomyocytes, the average cell catch per plate was 58 ± 4% (ranging from 22-84%). The average number of cells per plate with a seal resistance >0.25 GΩ was 48 ± 5% (range 11-80%). Among those cells, the percentage of cells expressing sodium current was 56 ± 4% (range 12-91%). Only cells with good voltage control and a peak current ≥300 pA were selected for analysis. Using these criteria, the success rate of recording sodium current from hiPSC-derived cardiomyocytes was 25 ± 2%. The average seal resistance of the cells selected for analysis was 0.618 ± 0.05 GΩ and the average capacitance was 18.3 ± 0.7 pF.

Drugs were diluted in the external solution and prepared in a separate 24 well plate. GS-967 was obtained from Gilead (Foster City, CA). Eleclazine, lidocaine, ranolazine and lacosamide were obtained from Sigma Aldrich (St. Louis, MO). DMSO concentration was the same for each concentration of a given drug. To block voltage-gated Ca^2+^ channels, 0.5 μM nisoldipine (Sigma Aldrich; St. Louis, MO). was added to external solutions.

### Data analysis

Cells were excluded from analysis if the maximum peak current was less than 300 pA. Patch-clamp measurements are presented as means ± SEM. Half maximal inhibitory concentration (IC_50_) values were calculated by fitting the dose response curves with a four parameter logistic equation where %_inhibition_ = Min_%inhibition_ + (Max_%inhibition_ – Min_%inhibition_)/(1 + ([drug]/IC_50_)^−n^), in which %_inhibition_ represents the percentage of peak current inhibited after block by each drug, Min_%inhibition_ and Max_%inhibition_ are the minimal and maximum % block observed for each drugs, [drug] is the concentration of drug, and n is the slope. The apparent binding rate (K_ON_) was measured using the time-dependent inhibition of I_Na_ during a variable length conditioning pulse. The apparent unbinding rate (K_OFF_) was measured using the time-dependent delay in the recovery of I_Na_ following an inactivating conditioning pulse.

## RESULTS

### Eleclazine and GS-967 exert limited tonic block in hiPSC-derived cardiomyocytes

We previously reported that 1 μM GS-967 inhibits 18% of *I*_*NaP*_ conducted by human Na_V_1.5 channels expressed in heterologous (tsA201) cells (Potet et al., 2016). Based on this finding we predicted that GS-967 and eleclazine would exhibit similar effects on *I*_*NaP*_ in human iPSC-derived cardiomyocytes. We performed automated patch clamp recordings of hiPSC-derived cardiomyocytes using the Syncropatch 768PE platform. Cardiomyocytes were held at −120 mV and currents were evoked by depolarizations to – 10 mV every 5 s. This infrequent pulsing protocol reveals drug interactions with the resting or open state of the channel and minimizes the effects of UDB, thereby providing a reasonable estimate of the extent to which GS-967 and eleclazine produce *I*_NaP_ tonic block. In all experiments, we used the single dose method to test the compounds. As shown in Fig. 1, GS-967 or eleclazine at 1 μM exhibited limited tonic block of *I*_NaP_ (14 ± 2% and 6 ± 3% for GS-967 and eleclazine respectively; n = 25-27). The maximum tonic block observed for GS-967 and eleclazine was 34 ± 3% and 43 ± 7% at 10 and 100 μM, respectively (Fig. 1). Higher concentrations could not be assayed due to the limited solubility of both compounds. To study the pharmacology of I_Na_ in hiPSC-derived cardiomyocytes we recorded sodium current from more than 2500 cells (Fig. S1).

**Figure 1.**
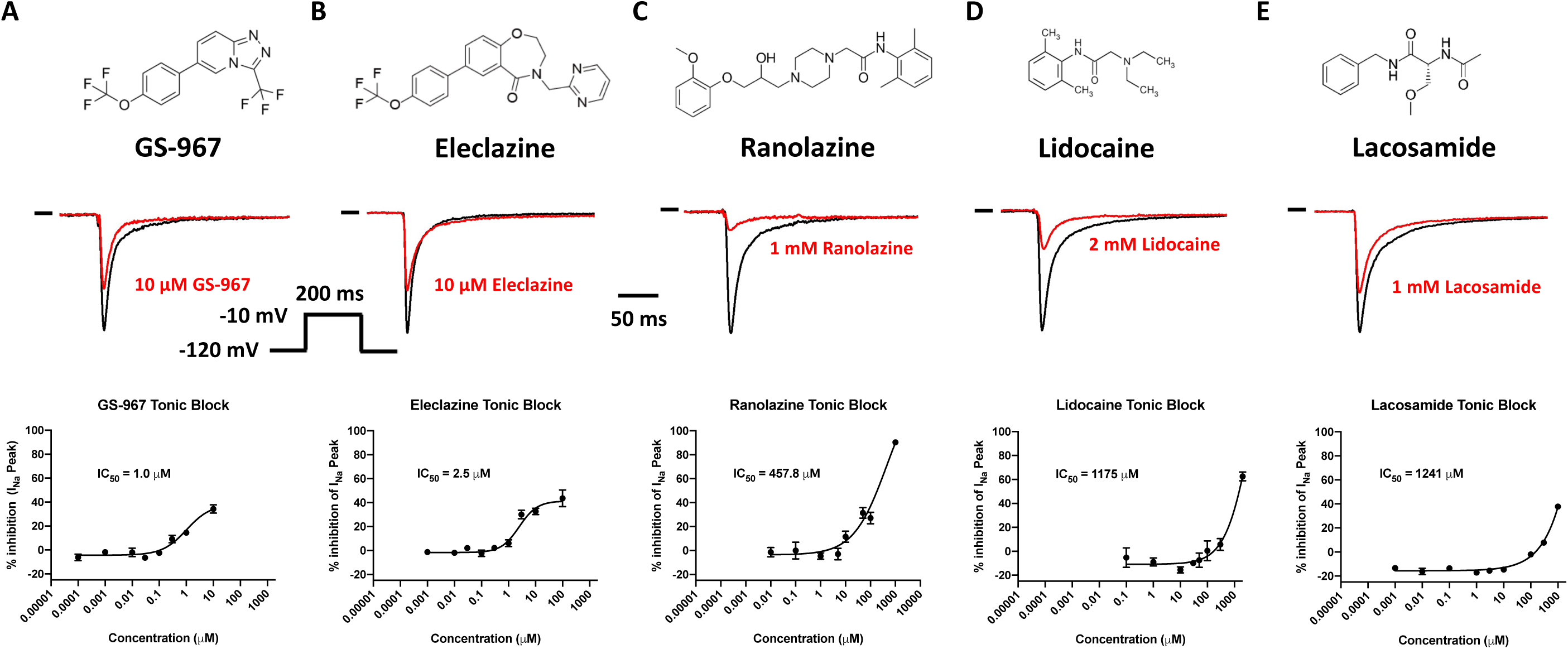
*I*_*NaP*_ tonic block in hiPSC-derived cardiomyocytes. Top: Drug chemical structures. Middle: representative *I*_NaP_ traces in absence (black) and presence (red) of the drug. Bottom: *I*_NaP_ concentration-response curve for GS-967 (**A**), eleclazine (**B**), ranolazine (**C**), lidocaine (**D**) and lacosamide (**E**). Currents were recorded from hiPSC-derived cardiomyocytes using automated patch clamp. Human iPSC-cardiomyocytes were held at a membrane potential of −120 mV and depolarized every 5 seconds to −10 mV for 200 ms (see inset). Peak sodium currents were measured at −10 mV before drug and were measured again 5 min after addition of the drug. The data are shown as a percentage of the peak current normalized to the maximum current in the absence of drug (*I/I*_*max*_). The representative current traces shown for the blocking effects of GS-967, eleclazine, ranolazine, lidocaine and lacosamide on *I*_*NaP*_ 5 min after perfusion were chosen at a concentration giving the maximum tonic block. The curves described by the solid lines were fitted by a four-parameter logistic equation. Each data symbol in the concentration response curves represent mean ± SEM for n = 7-67 cells. Estimated IC_50_ values are indicated for each concentration-response curve.

Figure 1 shows the concentration-response curves for *I*_NaP_ tonic block comparing GS-967, eleclazine, ranolazine, lidocaine and lacosamide. GS-967 and eleclazine exhibited similar IC_50_ values (1 and 2.5 μM, respectively) and had limited tonic block efficacy. By contrast, ranolazine, lidocaine and lacosamide exerted stronger maximum tonic block (for the maximal concentration tested) but with lower potency (IC_50_ = 457, 1175 and 1241 μM, respectively). We concluded that GS-967 and eleclazine exert less tonic block of *I*_NaP_ in hiPSC-derived cardiomyocytes compared to ranolazine, lidocaine and lacosamide.

### Use-dependent block of *I*_*NaP*_ in hiPSC-derived cardiomyocytes

We previously demonstrated that GS-967 exerts a strong UDB of Na_V_1.5 *I*_*NaP*_ in heterologous cells with greater potency than lidocaine and ranolazine (Potet et al., 2016). Others have shown that eleclazine can also exert UDB of Na_V_1.5 expressed in tsA201 cells (El-Bizri et al., 2018a; El-Bizri et al., 2018b; El-Bizri et al., 2018c). This motivated us to examine whether GS-967 and eleclazine exert similar effects in human cardiomyocytes.

We compared UDB by GS-967, eleclazine, ranolazine, lidocaine, and lacosamide in hiPSC-derived cardiomyocytes using two different protocols. First, we applied a series of 30 short (20 ms) depolarizing pulses (to −20 mV) at two frequencies (2 and 10 Hz). Second, we used longer (400 ms) depolarizing pulses (−20 mV) at a low frequency (2 Hz) to mimic the physiological cardiac cycle. In the absence of drugs, there was a ∼4 and ∼14% reduction of channel availability (frequency dependent inactivation) with the short pulse protocol at 2 and 10 Hz, respectively, and a ∼23% reduction with the longer pulse protocol (Fig. S2, S3 and S4). Following bath application of drugs, repetitive pulsing was associated with a progressive reduction of Na_V_1.5 *I*_NaP_ consistent with UDB and frequency dependent inactivation of the channel (Fig. S2, S3 and S4). To quantify the extent of UDB, we normalized the current amplitude measured after drug exposure by the current measured before drug application (Fig. 2A). The potency of each drug at producing UDB was compared by plotting the concentration-response curves (Fig. 2B, C and D).

**Figure 2.**
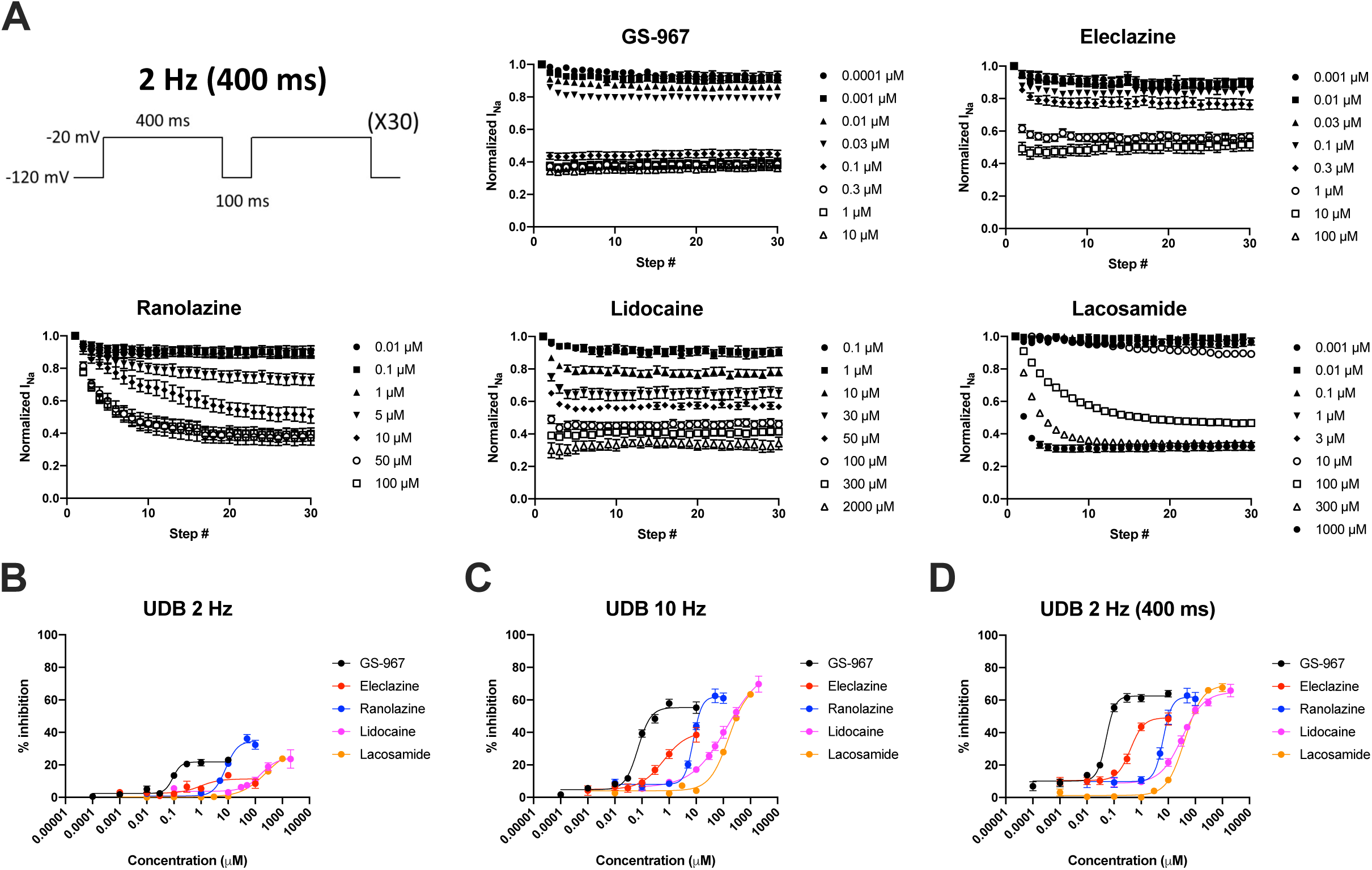
Concentration-response of *I*_*NaP*_ use-dependent block in hiPSC-derived cardiomyocytes by GS-967, eleclazine, ranolazine, lidocaine and lacosamide. **A**, To examine use-dependent block, hiPSC-derived cardiomyocytes were held at −120 mV and pulsed to −20 mV for 400 ms at 2 Hz, with an inter-pulse potential of −120 mV (see inset). The peak currents elicited by each pulse were normalized to the peak current of first pulse and plotted against the pulse number. The ratio after/before drug was plotted to assess the potency of peak *I*_*NaP*_ UDB. GS-967: concentrations from 0.0001 to 10 μM; n = 35-67. Eleclazine: concentrations from 0.0001 to 100 μM; n = 16-29. Ranolazine concentrations from 0.01 to 100 μM; n = 23-33. Lidocaine concentrations from 0.1 to 2000 μM; n = 6-27. Lacosamide concentrations from 0.001 to 1000 μM; n = 41-145. Concentration-response relationships for use-dependent block was studied using a series of 30 short (20 ms) depolarizing pulses (−20 mV) at two different frequencies (2 and 10 Hz) or longer physiological (400 ms) depolarizing pulses (−20 mV) at a low frequency (2 Hz). The % inhibition at the 30th pulse was calculated and plotted as a function of drug concentration. **B**, Concentration-response relationships for the use-dependent block at 2 Hz using a 20 ms step. IC_50_ = 0.09, 0.9, 8.1, 156.6 and 249.5 μM for GS-967, eleclazine, ranolazine, lidocaine and lacosamide respectively. **C**, Concentration-response for use-dependent block at 10 Hz using a 20 ms step. IC_50_ = 0.07, 0.6, 7.8, 133.5 and 158.5 μM for GS-967, eleclazine, ranolazine, lidocaine and lacosamide respectively. **D**, Concentration-response for use-dependent block at 2 Hz using a 400 ms step. IC_50_ = 0.05, 0.4, 6.5, 32.6 and 40.6 μM for GS-967, eleclazine, ranolazine, lidocaine and lacosamide respectively. All data are presented as mean ± SEM. The curves described by the solid lines were fitted by a four-parameter logistic equation.

All drugs exhibited limited UDB at 2 Hz using the 20 ms voltage step (Fig. 2B, Fig.S2). However, at 10 Hz and at 2 Hz using the longer protocol, all drugs showed greater UDB efficacy (Fig. 2C, 2D, Fig. S3 and S4). Eleclazine and GS-967 exhibited much greater UDB potency than ranolazine, lidocaine, and lacosamide. The order of UDB potency for the 5 drugs tested was the same as that found for tonic block. The calculated IC_50_ values for UDB at 10 Hz were 0.07 μM for GS-967, 0.6 μM for eleclazine, 7.8 μM for ranolazine, 133.5 μM for lidocaine and 158.5 μM for lacosamide (Figure 2C). At 10 Hz, eleclazine and GS-967 were 222-fold and 1907-fold more potent, respectively, than lidocaine, one of most well characterized use-dependent sodium channel blockers.

### Eleclazine and GS-967 affect recovery from inactivation and exhibit moderate unbinding kinetics

To explore plausible mechanisms to explain the UDB of *I*_*NaP*_ in hiPSC-derived cardiomyocytes by GS-967 and eleclazine, we first examined recovery from slow inactivation. This was done by utilizing a standard two-pulse protocol consisting of a depolarizing (−20 mV) 1000 ms pulse to reach maximum inhibition of peak I_Na_, followed by a variable duration recovery step to −120 mV and a final test pulse (−20 mV, 20 ms) (voltage protocol illustrated in Fig. S5). Channel availability after the end of the recovery interval was normalized to initial values and plotted against the recovery time. The time course of recovery was significantly slowed by all the drugs in a concentration-dependent manner (Fig. S5). The delayed recovery from inactivation reflects the dissociation of the drug from the channels, which can be quantified by comparing the recovery rate in the absence and presence of the compound (to remove drug-independent inactivation) (Fig.3). The unbinding rate (K_OFF_) was then calculated by fitting the dissociation time course. The calculated K_OFF_ was 1.6 s^−1^ and 1.5 s^−1^ for GS-967 and eleclazine, respectively, which is similar to lidocaine (1.1 s^−1^) but an order of magnitude slower than ranolazine (16.2 s^−1^) and 4-times more rapid than lacosamide (0.4 s^−1^) (Fig. 3).

**Figure 3.**
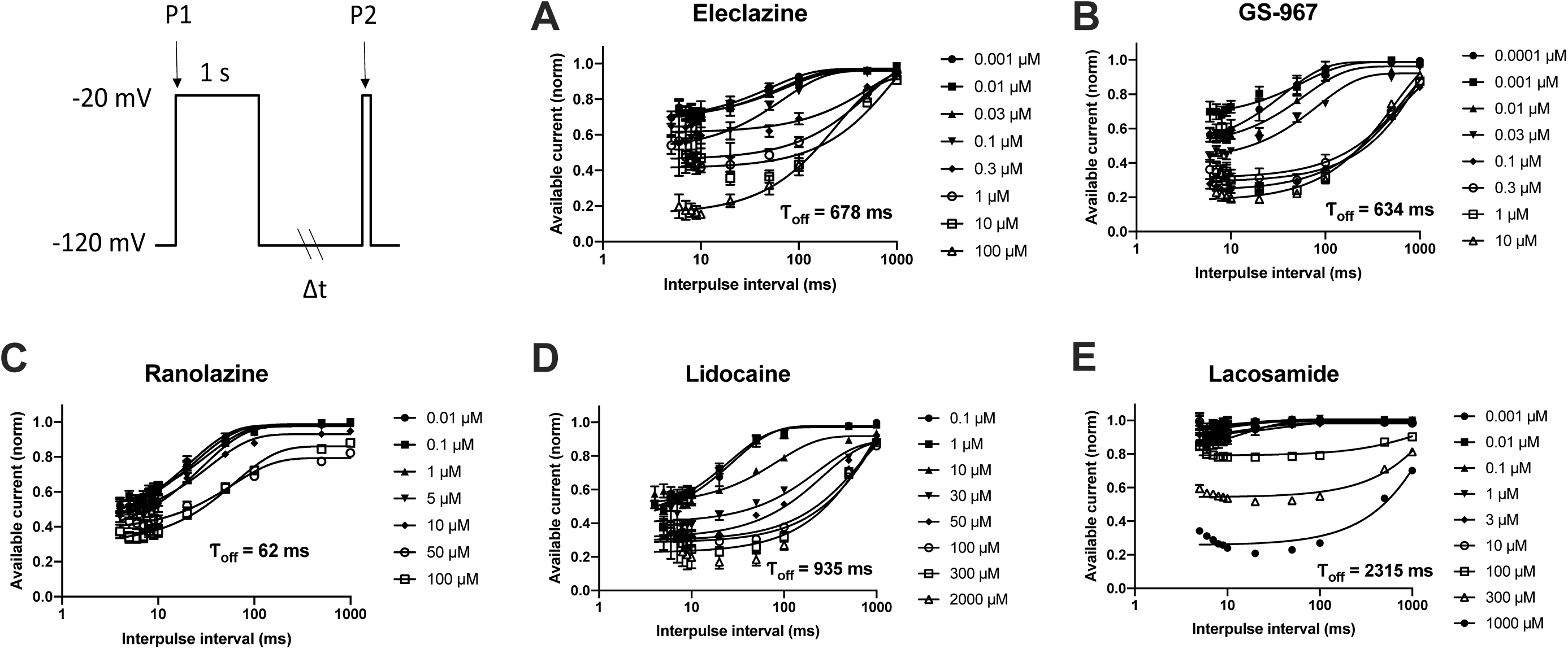
Recovery from inhibition in hiPSC-derived cardiomyocytes by GS-967, eleclazine, ranolazine, lidocaine and lacosamide. Recovery from inactivation was studied in hiPSC-derived cardiomyocytes with a standard two-pulse protocol consisting of a depolarizing (−20 mV) 1000 ms pulse to engage slow inactivation, followed by a variable duration recovery step to −120 mV and a final test pulse (−20 mV, 20 ms); see inset. Channel availability (recovery rate) after the end of the recovery interval was normalized to initial values and plotted against the recovery time. To calculate the unbinding rate (K_OFF_), the recovery of drug-bound channels was distinguished from non-bound channels by the ratio (I_DRUG_REC_/I_DRUG_MAX_)/(I_CTR_REC_/I_CTR_MAX_), where I_X_REC_ is the proportion of channels recovered in the presence or absence of a drug and I_X_MAX_ is the maximum current obtained in the presence or absence of drug (El-Bizri et al., 2018a). The recovery time course was fit with a single exponential equation to obtain a recovery time constant, τ_OFF_. The recovery time constant was averaged for each drug and the (K_OFF_) was calculated using K_OFF_ = 1/τ_OFF_: **A**, GS-967: K_OFF_ = 1.58 s^−1^; n = 12-32. **B**, eleclazine: K_OFF_ = 1.48 s^−1^; n = 23-24. **C**, ranolazine: K_OFF_ = 16.2 s^−1^; n = 16-19. **D**, lidocaine: K_OFF_ = 1.1 s^−1^; n = 3-10. **E**, lacosamide: K_OFF_ = 0.4 s^−1^; n = 73-127. Data represent mean ± SEM.

### Eleclazine and GS-967 exhibit rapid binding kinetics

Because UDB can be due to accumulation of channels in a slow inactivated state evoked by prolonged membrane depolarization, we investigated whether kinetic differences in the rate of entry into slow inactivation could account for UDB by GS-967 and eleclazine in hiPSC-cardiomyocytes. To assess the onset of slow inactivation, cells were held at −120 mV and then depolarized to −20 mV for a variable duration (2-1000 ms) followed by a brief recovery pulse (−120 mV for 20 ms) and a final 20 ms test pulse to −20 mV (voltage protocol illustrated in Fig. 4 and Fig. S6). The proportion of channels entering slow inactivation was estimated by normalizing to initial values (current recorded at P1) the current obtained after the short recovery pulse (current recorded at P2) and plotted against the first pulse duration. To assess the development of inhibition, we normalized the development of slow inactivation in the absence and presence of the compound to remove drug-independent inactivation (Fig. 4A-D). We could not measure the development of inhibition for ranolazine due its short dissociation time constant (τ_OFF_ = 62 ms, Fig. 3). The brief 20 ms recovery pulse to −120 mV was long enough to allow ranolazine to apparently unbind. The time-dependent inhibition of the Na^+^ currents shown in (Fig. 4A-D) was fitted with an exponential decay function having time constant τ. The 1/τ values were then plotted against the drug concentrations, and best fitted with a linear function (Fig. 4E). The slope represents the apparent binding rate (K_ON_) of the drug. As shown in Fig. 4E, the K_ON_ for GS-967, eleclazine, lidocaine and lacosamide were 25.7, 4.6, 0.2 and 0.003 μM^−1^.s^−1^, respectively. The K_ON_ for ranolazine was estimated from the Kd value (6.5 μM; equivalent to the K_OFF_/K_ON_ ratio). This Kd value is nearly identical to the IC_50_ value measured in the dose-response curve (Fig. 2D). The estimated K_ON_ for ranolazine was 2.5 μM^−1^.s^−1^. Data for all 5 drugs are summarized in Fig. 5. Eleclazine and GS-967 exhibit a higher association rate compared to ranolazine, lidocaine and lacosamide, but have rapid dissociation rates similar to lidocaine. Based on this analysis, the greater UDB potency of GS-967 and eleclazine is well correlated with the significantly higher association rates of these two compounds.

**Figure 4.**
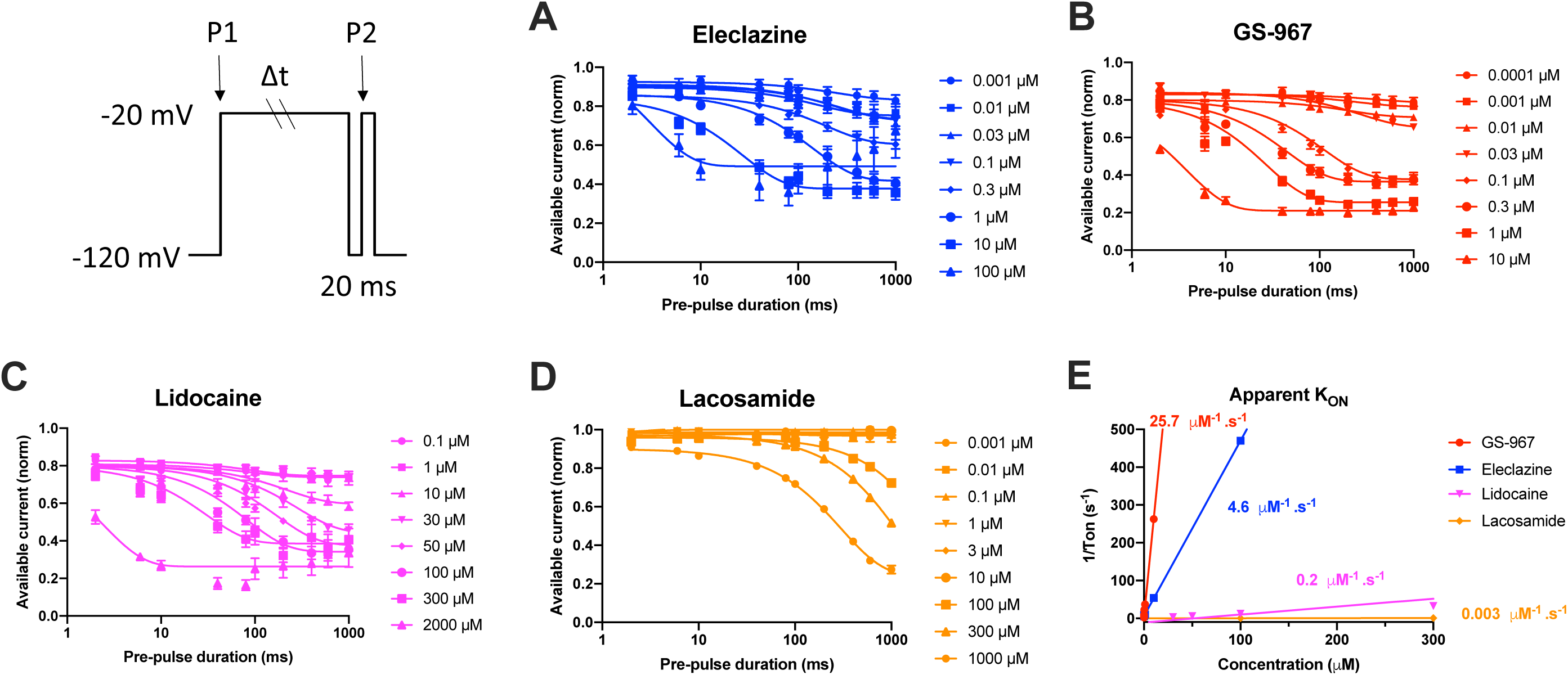
The apparent inhibition rate (K_ON_) of GS-967 and eleclazine is more rapid than ranolazine, lidocaine and lacosamide. The onset of slow inactivation induced by GS-967, eleclazine, ranolazine, lidocaine and lacosamide were determined in hiPSC-derived cardiomyocytes using the two-pulse protocol illustrated in the inset. Concentration-response curves were plotted using the data collected after a pre-pulse of 1000 ms. We normalized the development of slow inactivation in the absence and presence of the compound to remove drug-independent inactivation. The time-dependent inhibition of *I*_*NaP*_ shown for eleclazine (**A**), GS-967 (**B**), lidocaine (**C**), lacosamide (**D**) was fitted by an exponential decay function with a time constant (τ). **E**, Value for 1/τ were plotted against the drug concentrations, and best fitted by a linear function. The slope represents the apparent binding rate (K_ON_) of the drug. K_ON_ values for GS-967, eleclazine, lidocaine and lacosamide were 25.7, 4.6, 0.2 and 0.003 μM^−1^.s^−1^, respectively. Data represent mean ± SEM.

**Figure 5.**
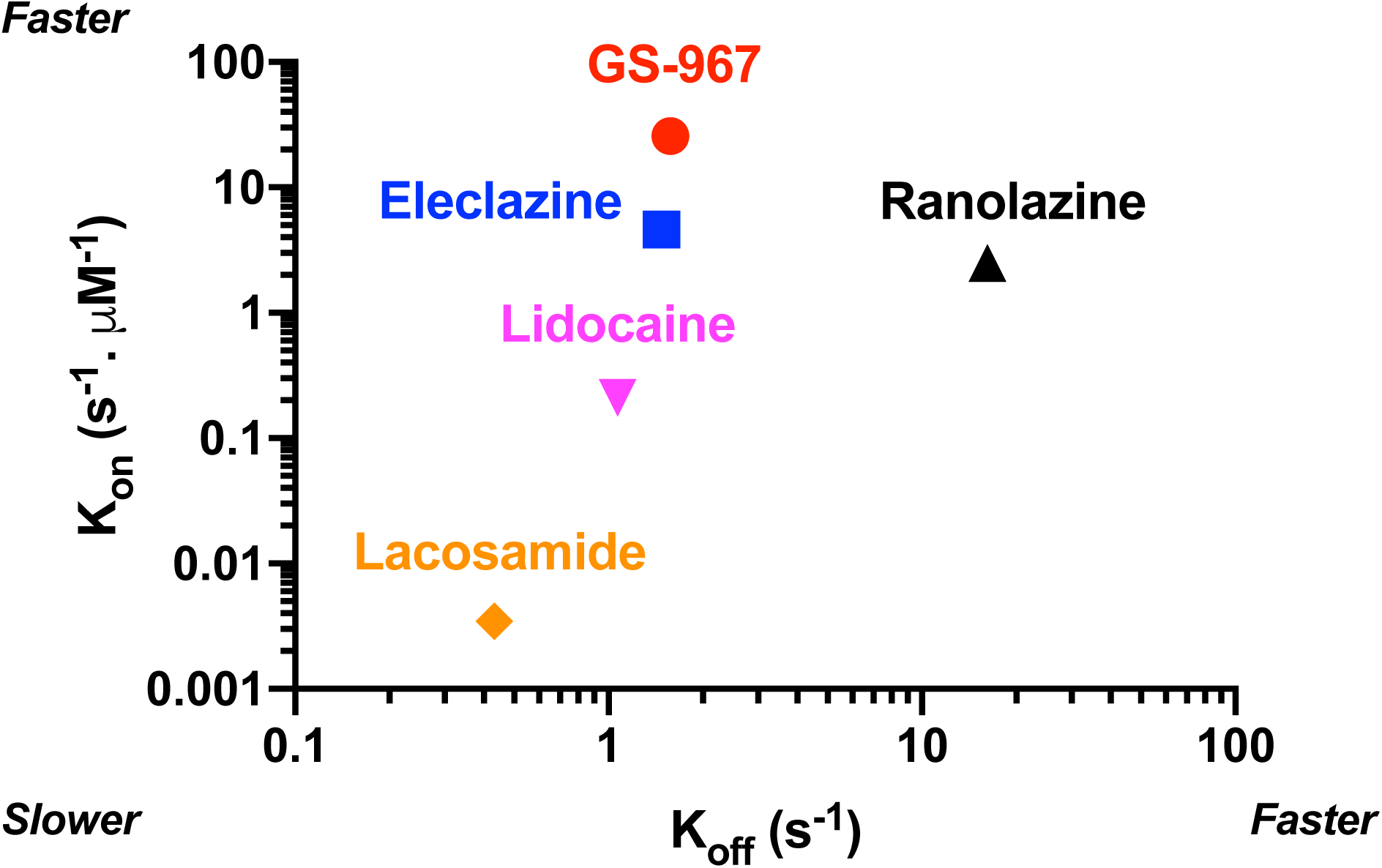
Relationship between K_ON_ and K_OFF_ for GS-967, eleclazine, ranolazine, lidocaine and lacosamide in hiPSC-derived cardiomyocytes. Relationship between the measured binding rate (K_ON_) and the measured unbinding rate (K_OFF_) for GS-967, eleclazine, lidocaine and lacosamide in hiPSC-derived cardiomyocytes. The K_ON_ for ranolazine was estimated as 6.5 μM from K_d_ = K_OFF_/K_ON_. This Kd value is nearly identical to the IC_50_ value measured in the dose-response curve (Fig. 2D).

## DISCUSSION

In this study, we used hiPSC-derived cardiomyocytes and automated patch clamp recording to elucidate the mechanism of action of two novel sodium channel blockers with demonstrated potent antiarrhythmic effects in various *in vitro* and *in vivo* models. Previously, the antiarrhythmic effects of GS-967 and eleclazine were attributed to preferential suppression of *I*_NaL_ (Bacic et al., 2017; Belardinelli et al., 2013; Burashnikov et al., 2015; El-Bizri et al., 2018a; Fuller et al., 2016; Pezhouman et al., 2014). More recently, it has been suggested that the antiarrhythmic effect of GS-967 was independent of the amplitude of *I*_NaL_ (Bossu et al., 2018). Here we offer evidence for other potentially important biophysical effects of GS-967 and eleclazine on sodium channels in human cardiomyocytes. We specifically demonstrated that GS-967 and eleclazine exert strong effects on slow inactivation and recovery from inactivation resulting in a substantial UDB similar to class 1-b antiarrhythmic drugs. These revelations may help explain the pharmacological effects of GS-967 and eleclazine in arrhythmia models.

A novel feature of our study is the use of automated planar patch clamp recording of hiPSC-derived cardiomyocytes in a 384-well configuration. While the use of planar patch clamp to record Na^+^ current in hiPSC-derived cardiomyocytes was described previously (Li et al., 2019; Rajamohan et al., 2016), these reports used much lower throughput platforms capable of recording from only 4-8 cells at a time. Our use of the 384-well platform in our study enabled a more robust capability to examine the pharmacological actions of multiple drugs simultaneously. The higher throughput also allowed us to use test single drug concentrations per well, and to measure binding and unbinding kinetics for 5 different drugs. These data demonstrate the feasibility of studying the pharmacological actions of drugs on the native human cardiac sodium current in hiPSC-derived cardiomyocytes.

Unique to our study, we demonstrated that GS-967 and eleclazine exert potent UDB of *I*_*NaP*_ in hiPSC-derived cardiomyocytes. Previously, UDB of GS-967 and eleclazine has not been examined in hiPSC-derived cardiomyocytes (Alves Bento et al., 2015; Belardinelli et al., 2013; Bonatti et al., 2014; Burashnikov et al., 2015; Carneiro et al., 2015; El-Bizri et al., 2018a; Fuller et al., 2016; Portero et al., 2017). Among the sodium channel blockers we tested, GS-967 exhibited the greatest UDB potency with IC_50_ values ranging from 50 to 90 nM depending on stimulation frequency and duration of the voltage steps (Fig. 2). GS-967 and eleclazine exerted UDB that was qualitatively similar to lidocaine (a prototypic use-dependent blocker of sodium channels) but with a significantly greater potency. The greater UDB potency was correlated with the unique on and off rate kinetics for these novel sodium channel blockers. By contrast, the rapid dissociation of ranolazine (K_OFF_, Fig. 3 and 5) and the slow binding kinetics of lacosamide (K_ON_, Fig. 4 and 5) were both correlated with less potent UDB.

GS-967 and eleclazine were originally demonstrated to exert potent antiarrhythmic effects in rabbit ventricular, canine and pig myocytes/heart by a proposed mechanism of action involving preferential *I*_NaL_ block (Bacic et al., 2017; Belardinelli et al., 2013; Fuller et al., 2016; Sicouri et al., 2013). In a more recent study, despite completely abolishing dofetilide-induced torsades de pointes (TdP) in a chronic atrioventricular block dog model, GS-967 did not completely suppress early afterdepolarizations (EADs) (Bossu et al., 2018). The authors concluded that GS-967 predominantly affects the perpetuation, but not the initiation, of arrhythmic events into TdP and that *I*_NaL_ density does not play a critical role in the moderate *in vitro* antiarrhythmic effect. Because the GS-967 and eleclazine concentrations used in those studies range from 0.2-1 μM, a range sufficient to suppress *I*_NaL_ and also to evoke UDB, we speculate that antiarrhythmic effects of these compounds may be due to the combination of *I*_NaL_ suppression and UDB preventing the perpetuation of the arrhythmia.

The effects of GS-967 and eleclazine resemble the effects of lidocaine, a class-Ib antiarrhythmic drug (Fig. 2). Effective class I antiarrhythmic drugs exhibit a greater efficacy in situations associated with rapid repetitive firing of action potentials (UDB) or prolonged tissue depolarization. Thus, effective arrhythmia suppression depends on the properties of the drug molecule that convey high affinity binding to the receptor on the channel pore when the channel is in the open or inactivated state. Such high affinity binding results in a slowed recovery of the drug-bound channel from inactivation as the cell membrane repolarizes (Ragsdale et al., 1996). In this study, we show that GS-967 and eleclazine, in hiPSC-derived cardiomyocytes, have very high association rates and moderate residence time comparable to lidocaine (Fig. 5). The moderate unbinding kinetics observed for GS-967 and eleclazine would limit peak I_Na_ inhibition and maintain the conduction velocity (Rajamani et al., 2016). The rapid binding of GS-967 and eleclazine would promote inhibition of late I_Na_ during phase 2 and 3 of the action potential and exert an antiarrhythmic action in the context of LQT3 syndrome (El-Bizri et al., 2018c). This was the rationale for clinical trials of eleclazine for type 3 LQTS (Gilead, 2000-2015).

In conclusion, we demonstrated the feasibility of using high throughput automated patch clamp recording to examine block of cardiac sodium current by multiple drugs in hiPSC-derived cardiomyocytes. We also demonstrated that GS-967 and eleclazine are more potent use-dependent blockers of cardiomyocyte sodium current than the antiarrhythmic drugs lidocaine and ranolazine or the antiepileptic drug lacosamide. We propose that potent UDB contributes to the antiarrhythmic effects of GS-967 and eleclazine.

## ACKNOWLEDGMENTS

The authors thank Hui-Hsuan Kuo for her technical assistance.

## AUTHORSHIP CONTRIBUTIONS

*Participated in research design:* Potet and George.

*Conducted experiments:* Potet and Egecioglu.

*Performed data analysis:* Potet.

*Wrote or contributed to the writing of the manuscript:* Potet, Burridge, and George.

## FOOTNOTES

This work was funded in part by a grant from Fondation Leducq and through research investments by the Northwestern Medicine Catalyst Fund.

## DISCLOSURES

Dr. Potet is a paid consultant for Praxis Precision Medicines, Inc. Dr. George serves on a scientific advisory board for Amgen, Inc. and received previous grant support from Merck and Co., Gilead Sciences, Inc., and Praxis Precision Medicines, Inc.

## Notes

### Competing Interest Statement

A.L.G. received a research grant from Praxis Precision Medicines, Inc for an unrelated research project involving one of the compounds studied in this manuscript (GS-967).

