## Supplemental Information for "GS-967 and Eleclazine Block Sodium Channels in Human Induced Pluripotent Stem Cell-derived Cardiomyocytes"

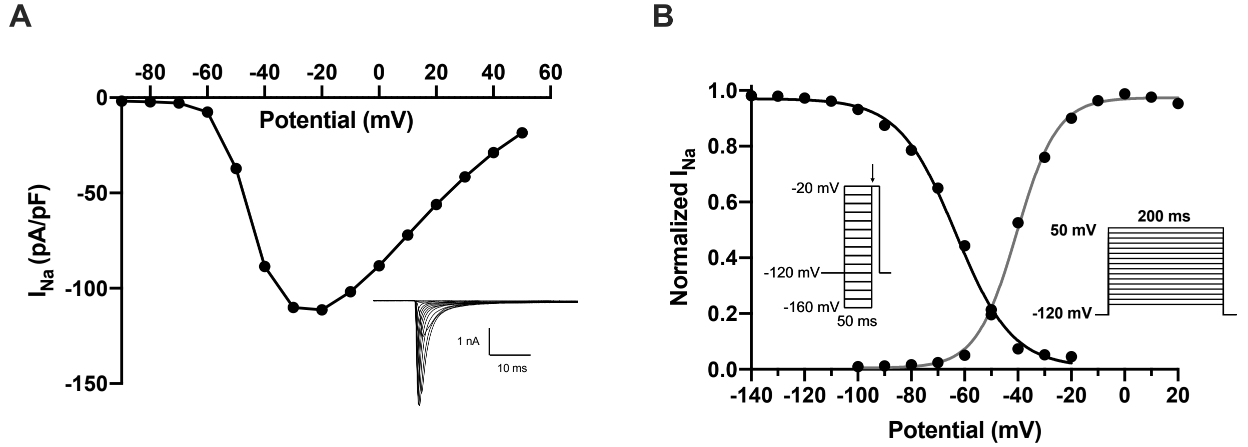


**Fig. S1. Biophysical properties of $I_{Na}$ in human iPSC-derived cardiomyocytes.**
**A,** averaged current-voltage relationship of $I_{Na}$ from hiPSC-derived cardiomyocytes (inset: averaged current traces). **B,** $I_{Na}$ voltage dependence of activation and inactivation in hiPSC-derived cardiomyocytes. Inset: voltage protocols used to assess voltage dependence of activation and inactivation. The half maximum voltage for inactivation was $V_{0.5}$ = -23.6 ± 0.1 mV (n = 2686). The half maximum voltage for activation was $V_{0.5}$ = -39.4 ± 0.2 mV (n = 1600). $V_{0.5}$ were determined by using the Boltzmann equation. All data are presented as mean ± SEM.

### SUPPLEMENTAL MATERIAL

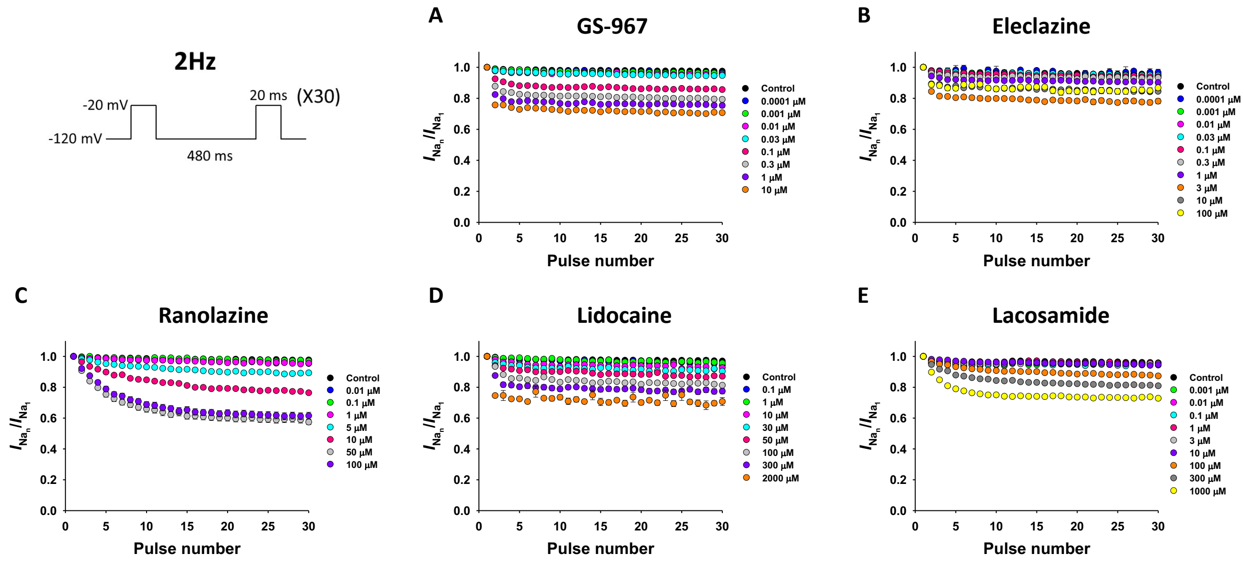


**Fig. S2. Use-dependent block of $I_{NaP}$ in human iPSC-derived cardiomyocytes.**
To examine use-dependent block, cells were held at -120 mV and pulsed to -20 mV for 20 ms at 2 Hz, with an inter-pulse potential of -120 mV (see inset). The peak currents elicited by each pulse were normalized to the peak current of first pulse and plotted against the pulse number. Black symbols represent absence of drug (control), while colored symbols represent experiments in the presence of different concentrations of the drugs tested. **A,** $I_{NaP}$ UDB response to GS-967 (0.0001 - 10 µM); n = 56-100. **B,** $I_{NaP}$ UDB response to eleclazine (0.0001 - 100 µM); n = 4-76. **C,** $I_{NaP}$ UDB response to ranolazine (0.01 - 1000 µM); n = 3-36. **D,** $I_{NaP}$ UDB response to lidocaine (0.1 - 2000 µM); n = 7-22. **E,** $I_{NaP}$ UDB response to lacosamide (0.001 to 1000 µM); n = 37-150. All data are presented as mean ± SEM.

### SUPPLEMENTAL MATERIAL

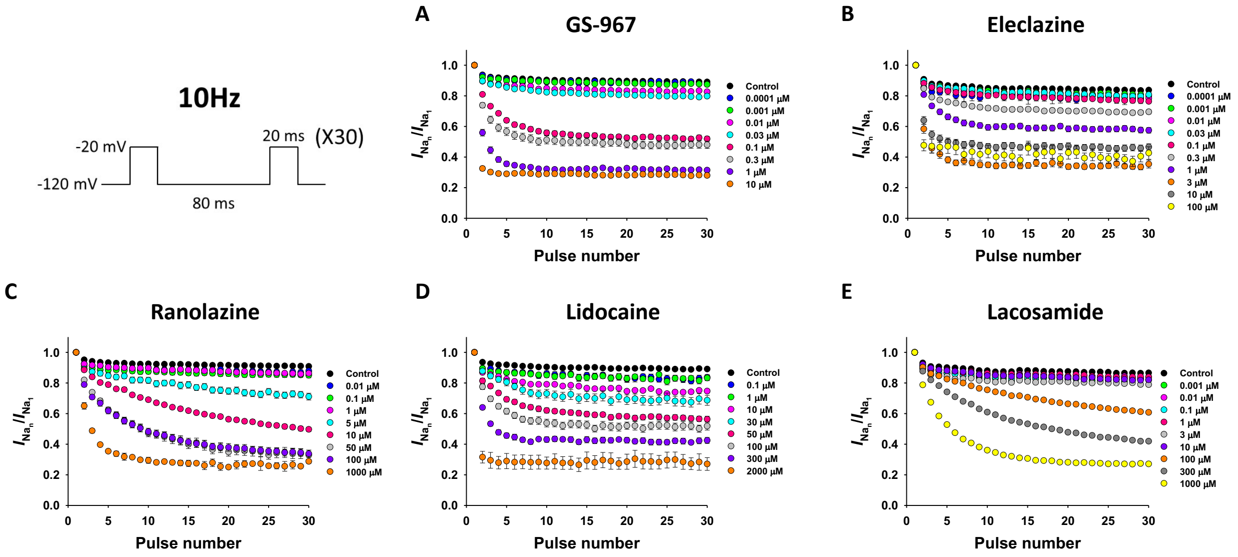


**Fig. S3. Use-dependent block of *I<sub>NaP</sub>* in human iPSC-derived cardiomyocytes.**
To examine use-dependent block, cells were held at -120 mV and pulsed to -20 mV for 20 ms at 10 Hz, with an inter-pulse potential of -120 mV (see inset). The peak currents elicited by each pulse were normalized to the peak current of first pulse and plotted against the pulse number. Black symbols represent absence of drug (control), while colored symbols represent experiments in the presence of different concentrations of the drugs tested. **A,** $I_{NaP}$ UDB response to GS-967 (0.0001 - 10 µM); n = 53-91. **B,** $I_{NaP}$ UDB response to eleclazine (0.0001 - 100 µM); n = 4-64. **C,** $I_{NaP}$ UDB response to ranolazine (0.01 - 1000 µM); n = 2-34. **D,** $I_{NaP}$ UDB response to lidocaine (0.1 - 2000 µM); n = 6-21. **E,** $I_{NaP}$ UDB response to lacosamide (0.001 - 1000 µM); n = 39-144. All data are presented as mean ± SEM.

### SUPPLEMENTAL MATERIAL

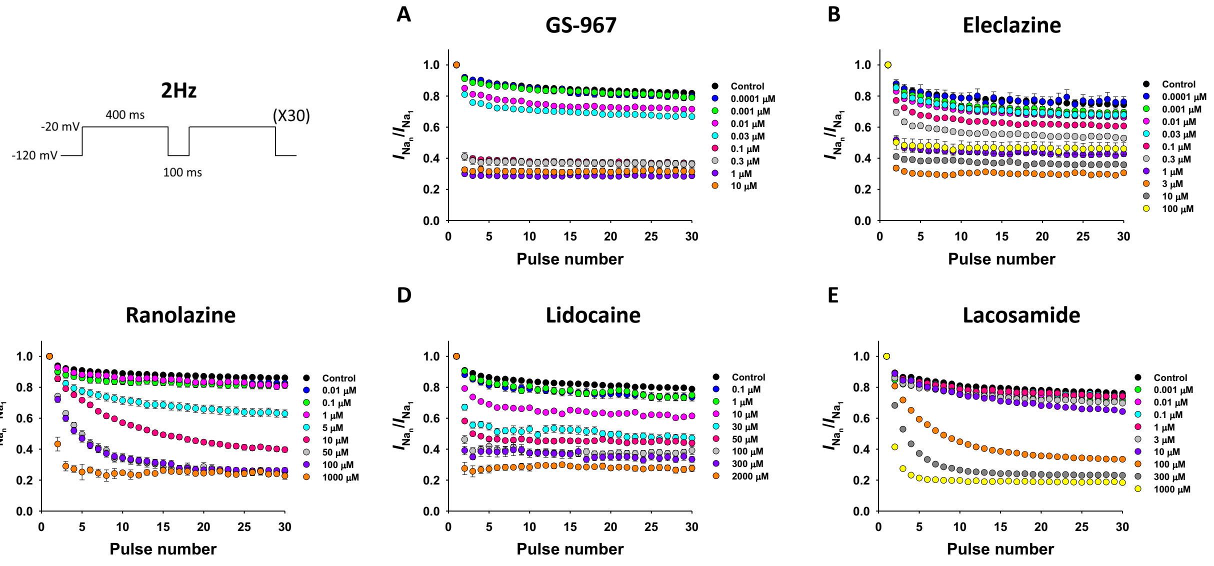


**Fig. S4. Use-dependent block of $I_{NaP}$ in human iPSC-derived cardiomyocytes.**
To examine use-dependent block, human iPSC-derived cardiomyocytes were held at -120 mV and pulsed to -20 mV for 400 ms at 2 Hz, with an inter-pulse potential of -120 mV (see inset). The peak currents elicited by each pulse were normalized to the peak current of first pulse and plotted against the pulse number. Black symbols represent absence of drug (control), while colored symbols represent experiments in the presence of different concentrations of the drugs tested. **A,** $I_{NaP}$ UDB response to GS-967 (0.0001 - 10 µM); n = 42-79. **B,** $I_{NaP}$ UDB response to eleclazine (0.0001 - 100 µM); n = 4-76. **C,** $I_{NaP}$ UDB response to ranolazine (0.01 - 1000 µM); n = 2-35. **D,** $I_{NaP}$ UDB response to lidocaine (0.1 - 2000 µM); n = 6-22. **E,** $I_{NaP}$ UDB response to lacosamide (0.001 -1000 µM); n = 38-145. All data are presented as mean ± SEM.

### SUPPLEMENTAL MATERIAL

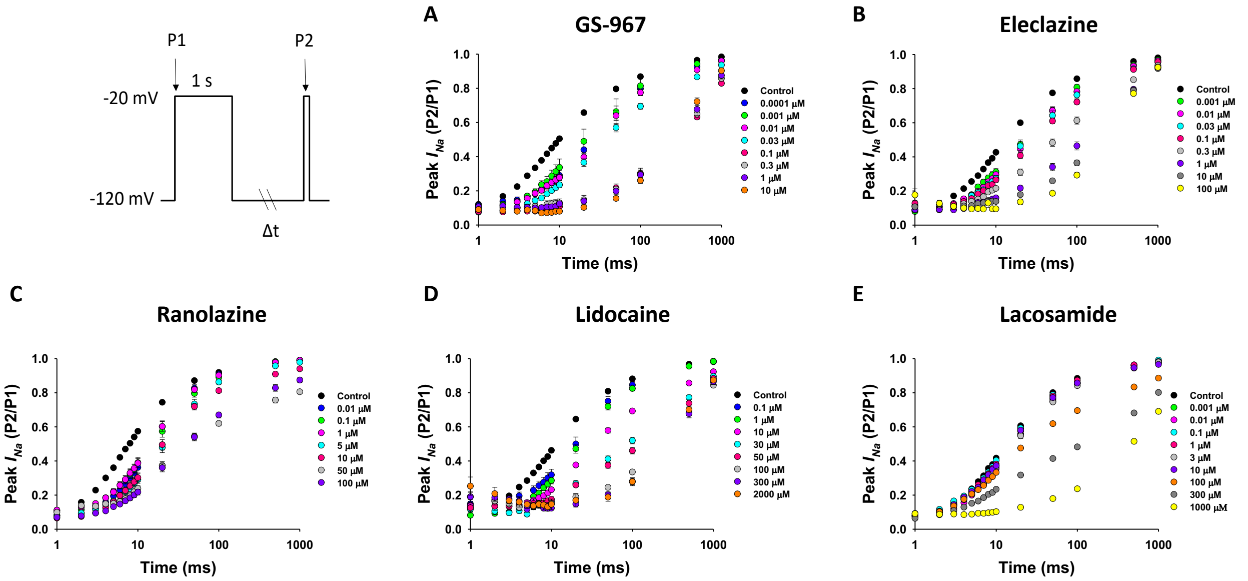


**Fig. S5. Effect of GS-967, eleclazine, ranolazine, lidocaine and lacosamide on recovery from inactivation in hiPSC-derived cardiomyocytes.**

Recovery from inactivation in hiPSC-derived cardiomyocytes was studied by utilizing a standard two-pulse protocol consisting of a depolarizing (-20 mV) 1000 ms pulse to engage slow inactivation, followed by a variable duration recovery step to -120 mV and a final test pulse (-20 mV, 20 ms); see inset. Channel availability after the end of the recovery interval (P2) was normalized to initial value (P1) and plotted against the recovery time. Recovery from inactivation was determined in the absence (black symbols) or presence of different concentrations (colored symbols) of GS-967, eleclazine, ranolazine, lidocaine and lacosamide. **A,** Recovery from inactivation response to GS-967 (0.0001 - 10 µM); n = 6-27. **B,** Recovery from inactivation response to eleclazine (0.001 - 100 µM); n = 25-40. **C** Recovery from inactivation response to ranolazine (0.01 - 1000 µM); n = 14-30. **D,** Recovery from inactivation response to lidocaine (0.1 - 2000 µM); n = 5-17. **E,** Recovery from inactivation response to lacosamide (0.001 -1000 µM); n = 33-127. All data are presented as mean ± SEM.

### SUPPLEMENTAL MATERIAL

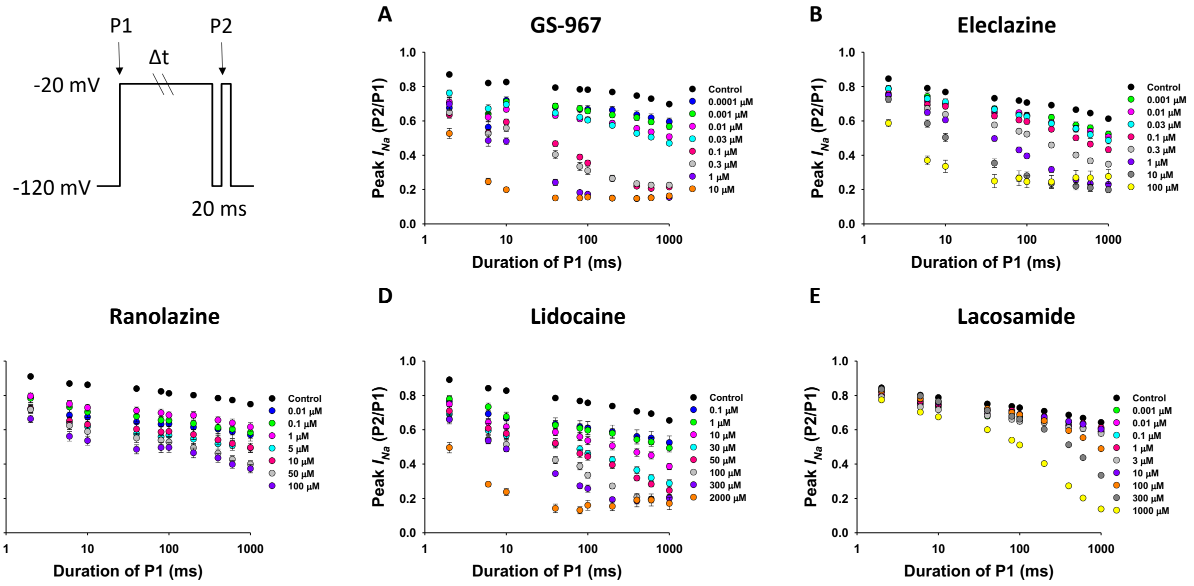


**Fig. S6. Effect of GS-967, eleclazine, ranolazine, lidocaine and lacosamide on onset of slow inactivation in hiPSC-derived cardiomyocytes.**

Onset of slow inactivation was determined in the absence (black symbols) and presence of different drug concentrations (colored symbols) using a two-pulse protocol (inset): hiPSC-derived cardiomyocytes were held at -120 mV and then depolarized to -20 mV for a variable duration (2-1000 ms) followed by a brief recovery pulse (-120 mV for 20 ms) and a final 20 ms test pulse to -20 mV. Channel entering slow inactivation were estimated by normalizing to initial values (current recorded at P1) the current obtained after the short recovery pulse (current recorded at P2) and plotted against the first pulse duration. **A,** Onset of slow inactivation response to GS-967 (0.001 - 10 μM); n = 11-84. **B,** Onset of slow inactivation response to eleclazine (0.001 - 100 μM); n = 33-72. **C,** Onset of slow inactivation response to ranolazine (0.01 - 1000 μM); n = 5-33. **D,** Onset of slow inactivation response to lidocaine (0.1 - 2000 μM); n = 6-22. **E,** Onset of slow inactivation response to lacosamide (0.001 - 1000 μM); n = 38-140. All data are presented as mean ± SEM.
